# Base Editing for Reprogramming Cyanobacterium *Synechococcus elongatus*

**DOI:** 10.1101/2022.09.05.506134

**Authors:** Shu-Yan Wang, Xin Li, Shu-Guang Wang, Peng-Fei Xia

**Affiliations:** School of Environmental Science and Engineering, Shandong University, Qingdao 266237, China; Sino-French Research Institute for Ecology and Environment, Shandong University, Qingdao 266237, China

**Keywords:** cyanobacteria, base editing, CRISPR, multiplex, *Synechococcus elongatus*

## Abstract

Global climate change demands carbon-negative innovations to reduce the concentration of atmospheric carbon dioxide (CO_2_). Cyanobacteria can fix CO_2_ from the atmosphere and can be genetically reprogrammed for the production of biofuels, chemicals and food products, making an ideal microbial chassis for carbon-negative biotechnology. However, the progress seems to be slowed down due to the lagging-behind synthetic biology toolkits, especially the CRISPR-Cas-based genome-editing tools. As such, we developed a base-editing tool based on the CRISPR-Cas system and deamination for cyanobacterium *Synechococcus elongatus*. We achieved efficient and precise genome editing at a single-nucleotide resolution, and identified the pure population of edited cells at the first round of selection without extra segregation. By using the base-editing tool, we successfully manipulated the glycogen metabolic pathway via the introduction of premature STOP codons to inactivate the corresponding genes. We demonstrated multiplex base editing by editing two genes at once, obtaining a nearly two-fold increase in the glycogen content. We present here the first report of base editing in the phylum of cyanobacteria, and a paradigm for applying CRISPR-Cas systems in bacteria. We believe that this work will accelerate the synthetic biology of cyanobacteria and drive more innovations to alleviate global climate change.

## Introduction

Bio-based economy alleviates the dependency on fossil-based fuels and chemicals, marking a milestone toward a sustainable future (1). As an important sector of bio-based economy, carbon-negative biotechnology deploys microorganisms to convert CO_2_ from either industrial waste gases or directly from the atmosphere to desired bioproducts (2, 3). Cyanobacteria use CO_2_ and solar energy for metabolism, and, as reported, they fix 10 – 20% of the global organic carbon sources (4), making them preferred photosynthetic chassis for the development of carbon-negative biotechnology.

Cyanobacteria are genetically amendable, and some of the strains, e.g., *Synechococcus elongatus* PCC 7942 and *Synechocystis* sp. PCC 6803, are naturally competent and with detailed genetic information (5, 6). Given the advances in metabolic engineering and synthetic biology, cyanobacteria have been engineered to produce a great variety of products by rewiring the metabolic networks (7, 8). For instance, regulating the metabolic pathway of glycogen can optimize the synthesis of target chemicals, such as sucrose, succinate, isobutanol, 1-butanol or glycogen per se (9-13). Despite the progress thereof, current tools for genome-editing in cyanobacteria are often labor-intensive and time-consuming, and the knock-in/out genome-editing may be challenging due to the polyploid chromosomes (14). Hence, novel genome-editing tools are necessary to explore the untapped potential of cyanobacteria.

CRISPR-Cas systems, the bacterial and archaeal immune systems, have been repurposed as a powerful genome-editing tool, and they have been promoting the development of biotherapeutics, biodiagnoses and bioproduction (15-17). To edit the genome of a bacterium, the CRISPR-Cas system targets a specific site navigated by a programmed guide RNA (gRNA) and introduces a double-strand break (DSB) at the designed locus, resulting in cell death or forcing the homology-directed recombination to repair the DSB with a donor DNA. Then, the genome will be edited without a marker or scar. As the CRISPR-Cas system cleaves all target sequences, it makes the single-round selection of edited microbes possible, and it may bypass the time-consuming segregation step in cyanobacteria (18, 19). Due to these superiorities, scientists have been trying to adapt CRISPR-Cas-based genome-editing in cyanobacteria (11, 20, 21). Li et al. (2016) first applied the CRISPR-Cas9 system from *Streptococcus pyogenes* to engineer *S. elongatus* PCC 7942 for improved succinate production. Whereas they have to utilize a transient expression system to lower the toxicity, which is a common observation in cyanobacteria (14, 20). This obstacled the development of CRISPR-Cas-based genome editing in cyanobacteria. Current progress of CRISPR-Cas systems in cyanobacteria mainly owes to CRISPR interference (CRISPRi), where nuclease deactivated Cas9 (dCas9), as the working effector, binds to the target DNA instead of cleaving it and shows much lower toxicity (5, 22, 23). Despite the availability of CRISPRi, complementary genome-editing methods are still in demand for genetic modification in cyanobacteria.

Base editing is a rising CRISPR-Cas-based genome-editing tool at a single-nucleotide resolution using dCas9 for targeting and deaminase for editing. It can generate cytosine-to-thymine (C-to-T) or adenine-to-guanine (A-to-G) substitutions without requiring donor DNAs or causing DSBs (24-26). For the Target-AID system, the dCas9 will target the protospacer located by the protospacer adjacent motif (PAM) under the guide of a programmed gRNA, and forms an R-loop. In the editing window, normally among the positions -16 to -19 upstream of the PAM, the cytidine deaminase will convert cytidine to uracil in the single-strand DNA of the R-loop, which will be read as thymine in the DNA repair process, generating C-to-T substitutions (27). By designing gRNAs, the gene of interest can be inactivated via the introduction of a premature stop codon (e.g., changing

CAA to TAA), which may result in the reprogramming of correlated metabolic pathways. Base editing has been successfully adapted in bacteria, such as *Streptomyces griseofuscus, Agrobacterium rhizogenes, Pseudomonas putida* and *Clostridium ljungdahlii* (28-31). To be noticed, Xia et al. (2020) reported the potential of base editing in metabolic engineering of *Clostridium ljungdahlii* via a genome-scale interrogation, and demonstrated that base editing could overcome the challenges of applying CRISPR-Cas-based genome-editing in bacteria, including the toxicity. Therefore, base editing may also shed light on the development of genome-editing tools for cyanobacteria.

Here, we developed a base-editing system for the model cyanobacterium *S. elongatus* PCC 7942. A modularized assembly procedure was devised to generate base-editing plasmids, and the C-to-T nucleotide substitutions were successfully demonstrated on the designed loci. We deployed the base-editing system to reprogram the glycogen metabolism in *S. elongatus* PCC 7942 by introducing premature STOP codons to inactivate corresponding genes. We also achieved multiplex base editing in two target genes at once, and significantly increased glycogen content was observed. As the first report of base editing in the phylum of cyanobacteria, our work provides a paradigm for applying CRISPR-Cas systems to promote carbon-negative biotechnology.

## Results

### Design and demonstration of base editing in *S. elongatus* PCC 7942

To construct the base-editing tool for *S. elongatus* PCC 7942, we first combined the dCas9 from *S. pyogenes* with the activation-induced cytidine deaminase (PmCDA1) from sea lamprey (*Petromyzon marinus*) to generate the editing module (27), and the uracil DNA glycosylase inhibitor (UGI) with a Leu-Val-Ala (LVA) degradation tag was employed to improve the editing efficiency and reduce the potential toxicity (32). We chose the inducible *lacI*-P_trc_ system (33) and the self-replicating plasmid pAM4787 (34) to drive and carry the editing module, successfully building the plasmid pSY. Then, the gRNAs targeting different loci were generated via inverse PCR on pTemplate. Finally, the two modules were assembled to obtain a “third-party” genome-editing system (Fig. 1A).

**Fig. 1.**
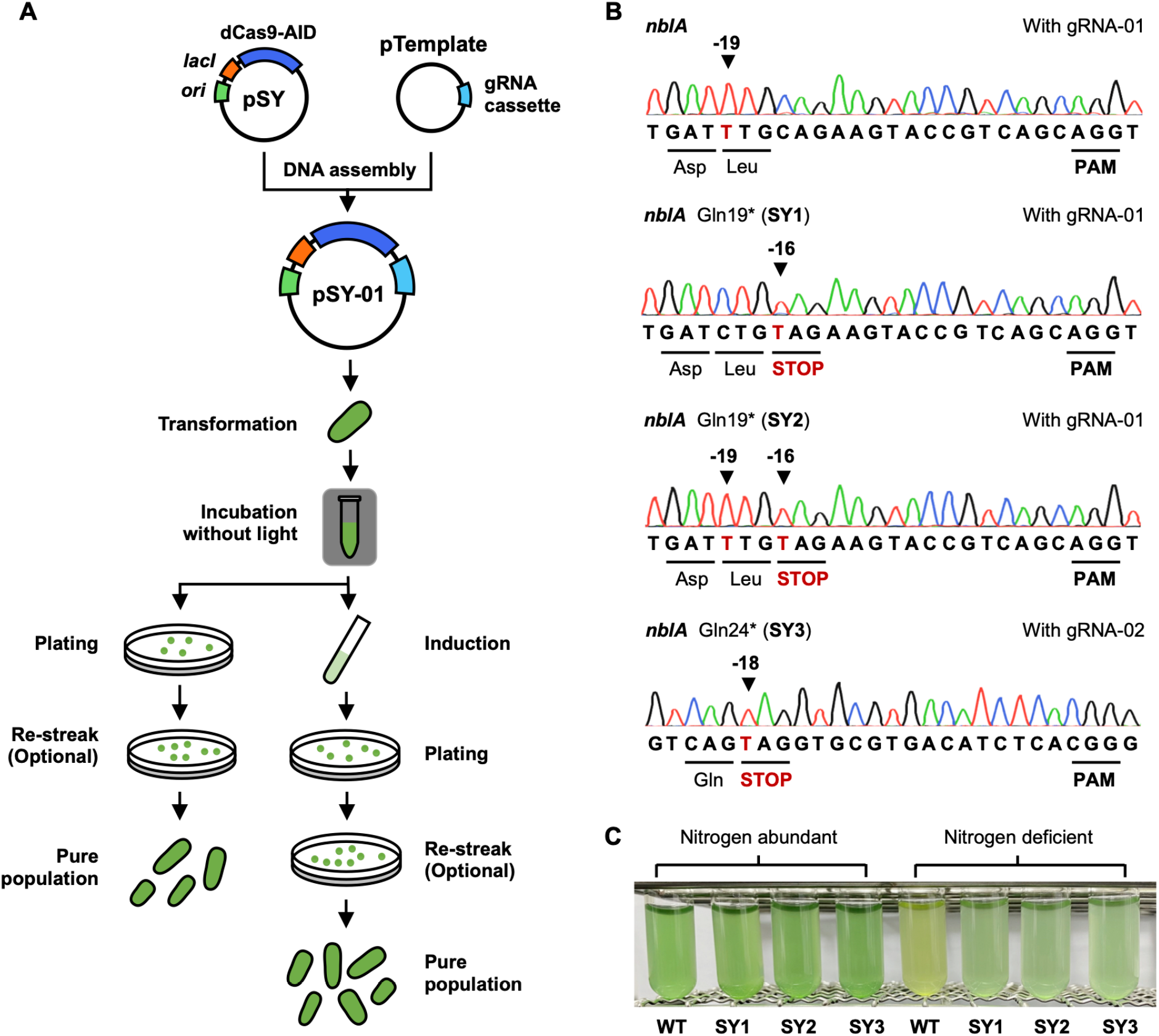
Base editing in cyanobacteria. **(A)** The workflow of base editing. First, the specific gRNA cassette was integrated into the plasmid pSY, containing the inducible *lacI*-P_trc_ system, the dCas9-AID cassette, and the gRNA cassette. Then, it was transformed into *S. elongatus* PCC 7942. Two induction strategies were tested. For the direct-induction method, the cells were induced on the selective plates, and for the liquid-induction method, the cells were induced in the recovery liquid medium before plating. The transformants were randomly picked for sequencing to check the editing. When required, the transformants were re-streaked to obtain the pure population of designed editing. **(B)** The sequencing results of the target loci in *nblA* after being edited with pSY-01 (gRNA-01) and pSY-02 (gRNA-02). The edited loci were highlighted by the black arrow and the positions were counted from the PAM, regarding the first base on the left side of a PAM as position -1. **(C)** Phenotypical examination of the edited strains. The *nblA*-inactivated strains (SY1, SY2 and SY3) and the wild-type strain were cultivated in BG-11 medium with and without nitrogen source for 48 h to check the bleaching phenotype resulted from *nblA*.

To demonstrate our base-editing system, we selected *nblA*, which encodes the phycobilisome hydrolyzing enzyme, as our target (35). By disrupting *nblA*, the strain will not display the bleaching phenotype (color change from green to yellow) in nitrogen starvation conditions. We designed gRNA-01 (Table S1) and the corresponding plasmid pSY-01, which may generate a C-to-T substitution at position -16 (the first base upstream of PAM as position -1) and introduce a premature STOP codon to inactivate *nblA*. After being transformed with pSY-01, the cells were induced directly on selective plates, and the resulting colonies were then randomly picked and sequenced. We selected 38 colonies in total from 3 independent rounds of editing, and observed an overall editing efficiency of 73.41% (Table S2). The editing included C-to-T substitutions at positions -19, -16, and both (Fig. 1B), which was in agreement with the previously reported editing window between positions -16 and -19 (27). In addition, we designed gRNA-02 (Table S1) and applied the corresponding plasmid pSY-02 to further demonstrate our system, and we observed precise editing in position -18 of the target sequence and introduced a premature STOP codon to *nblA* (Fig. 1B), reaching an editing efficiency of 27.27%. We tested the phenotype of three base-edited *S. elongatus* strains thereof, SY1 (*nblA* Gln19*, obtained with pSY-01), SY2 (*nblA* Gln19* and a silent bystander mutation at Leu18, obtained with pSY-01), and SY3 (*nblA* Gln24*, obtained with pSY-02) (Fig. 1B). All three edited strains harbored premature stop codon in *nblA*, leading to its inactivation. We cultivated these strains and the wild-type strain with and without nitrogen source for 48 h. As expected, the edited strains did not exhibit obvious variations in color while the wild-type strain showed an evidently bleaching phenotype in the nitrogen-starvation condition (Fig. 1C), revealing the success of base editing.

One common issue for base editing is the edited loci sometimes show mixed sequencing signals (29, 36), and the polyploid cyanobacterial chromosomes may make it even harder to get clean editing. We also found mixed sequencing signals in some of the edited loci, but to be noticed, we obtained clean editing immediately at the first round of selection, and the efficiency of clean editing reached 21.03% for gRNA-01 and 9.09% for gRNA-02 which accounted for 27.19% and 33.33% of all the editing, respectively (Table S2). This will considerably reduce the time for genome editing in cyanobacteria, where two rounds of segregation are normally required. The clean edited colonies were further verified to be pure populations after another round of segregation (Fig. S1A). At last, we demonstrated that a pure population with designed clean editing could be obtained from colonies with mixed sequencing signals after one more round of segregation (Fig. S1B).

Next step, we tried to elevate the efficiency and attempted a second induction method by inducing the cells in the liquid medium immediately following the 24-h incubation after the transformation of pSY-01. The induced cells were then plated on selective plates without inducers. The sequencing results showed a higher editing efficiency of 86.51% (p =0.224), 31.79% (p =0.699) of which are clean editing (Table S2, Fig. S2A and S2B). Though no statistically significant increase in efficiency was confirmed, we chose the liquid-induction approach in the following study to obtain higher editing efficiency. For the final step of genome editing, we found that the working plasmids could be easily cured in only one passage of cultivation in non-selective medium, and no extra genetic design, such as *sacB*-based suicide module, is required. This may result from the relatively low stability of the pAM4787 backbone (34), which makes it an ideal vector to carry genome editing tools for cyanobacteria.

### Reprograming glycogen metabolic pathway with base editing

We chose the glycogen metabolic pathway to examine whether the base-editing tool could reprogram the metabolism in cyanobacteria. Glycogen is a natural carbon sink and energy reservation compound for cyanobacteria, and it is a platform chemical for producing diverse bioproducts (9). For the synthesis of glycogen, CO_2_ is converted into glucose-1-phosphate through the Calvin cycle and enters the glycogen metabolic pathway. Catalyzed by glucose-1-phosphate adenylyltransferase (encoded by *glgC*) and glycogen synthase (encoded by *glgA*), glucose-1-phosphate is assembled into polyglucose chains, forming glycogen (Fig. 2A). For the degradation of glycogen, glycogen phosphorylase (encoded by *glgP*) and glycogen debranching enzyme (encoded by *glgX*) will hydrolyze glycogen, breaking it down and generating glucose-1-phosphate back or branched glucans (Fig. 2A) (4). Therefore, we aimed to inactivate the *glgP* and *glgX*, respectively, to increase the glycogen accumulation by cutting off one of the glycogen degradation pathways, and *glgC* was targeted to block the glycogen synthesis and to redivert the carbon flux.

**Fig. 2.**
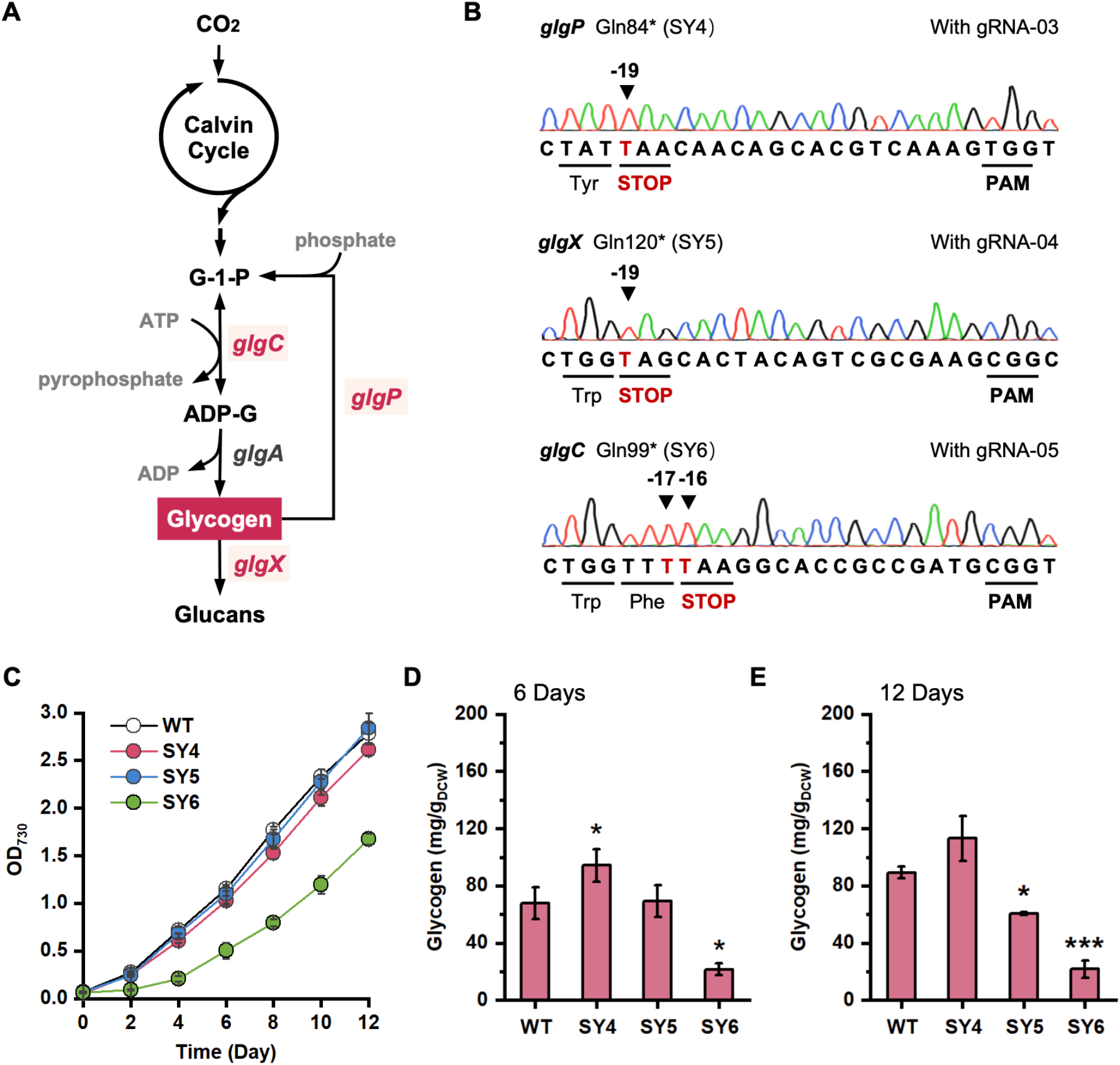
Reprogramming the glycogen metabolic pathways with base editing. **(A)** The glycogen metabolic network in cyanobacteria. **(B)** The sequencing results of the genes *glgP, glgX* and *glgC* in the edited strains (SY4, SY5, SY6). The edited loci were highlighted by the black arrow and the positions were counted from the PAM, regarding the first base on the left side of a PAM as position -1. **(C)** Growth profiles for the edited strains and the wild-type strain via the measurement of OD_730_ every 24 h. **(D)** Glycogen contents measured on day 6. **(E)** Glycogen contents measured on day 12. Glycogen was extracted and hydrolyzed into glucose before being analyzed by HPLC. The error bars represent the standard deviations of three independent experiments. The differences in glycogen contents between the edited strains and the wild type were verified by *t*-test (* for P < 0.05, ** for P < 0.01, and *** for P < 0.001).

We designed three base-editing plasmids, pSY-03, pSY-04 and pSY-05, with gRNA-03, gRNA-04, and gRNA-05 (Table S1), to introduce premature stop codons in *glgP, glgX*, and *glgC* for gene inactivation. By applying the three working plasmids, we successfully constructed the desired strains. Strain SY4 with inactivated *glgP* was obtained via generating a C-to-T substitution to change Gln84 to a stop codon, and notably, the strain was immediately identified at the first round of selection without extra segregation (Fig. 2B). For strains SY5 and SY6, we found mixed sequencing signals in all picked colonies at position -19 and -16 in the protospacers targeted by gRNA-04 and gRNA-05, respectively, while we observed bystander editing at position -16 for gRNA-04 and position -17 for gRNA-05 (Fig. S3B and S3C). After one more round of segregation, pure populations of SY5 with inactivated *glgX* (Gln120*) and of SY6 with inactivated *glgC* (Gln99*) were obtained (Fig. 2B).

Then, we verified the phenotypes of the three edited strains. We cultivated SY4, SY5 and SY6 with the wild-type strain and evaluated the accumulated glycogen contents. We observed that SY4 and SY5 showed similar growth profiles to that of the wild-type strain, while SY6 displayed a repressed growth, implying that the inhibition of glycogen synthesis, which was caused by inactivated *glgC*, may influence the growth of cyanobacteria (Fig. 2C). On day 6, SY4 showed a higher glycogen content of 94.28 ± 11.32 mg/gDCW compared with that of the wild-type strain (67.86 ± 11.29 mg/gDCW), and, as expected, SY6 showed a lower glycogen content with 21.64 ± 4.28 mg/gDCW (Fig. 2D). Given these results, the inactivation of *glgP* improved the glycogen accumulation by 38.93% (p = 0.0458), and the lost function of *glgC* reduced glycogen accumulation by 68.11% (p = 0.0112). These results correspond to previous reports that the deletion of *glgP* increases the glycogen content while the knockout of *glgC* leads to lower or trace amounts of glycogen (4). SY5 showed a similar glycogen content (66.22 ± 9.38 mg/gDCW, p =0.856) to the wild-type strain (Fig. 2D), indicating that the inactivation of *glgX* would not lead to glycogen accumulation. This was in agreement with the previous research which revealed that the loss of *glgX* would cause instability in glycogen content instead of its over-accumulation (4). On day 12, we observed a similar tendency for SY4 and SY6. Compared with the glycogen content of the wild-type strain, SY4 showed an increase of 26.70% (p = 0.262), while SY6 displayed a reduction of 75.71% (p = 0.0008). To our surprise, the glycogen content in SY5 decreased by 32.01% (p = 0.0537), while it remained similar on day 6 (Fig. 2E). These results demonstrated that base editing can reprogram the metabolism in cyanobacteria via a single-nucleotide substitution, while they also indicated that manipulating a single gene may not be sufficient for metabolism reprogramming.

### Multiplex base editing with tandem gRNAs

Editing multiple genes is normally required in reprogramming a microbe, but it consumes lots of time and labor since multiplex genome editing at one time is quite challenging in cyanobacteria. To our knowledge, only one recent study reported multiplex genome editing in cyanobacterium *Synechocystis* sp. PCC 6803 (37), and no such progress in *S. elongatus* have been documented. We explored the potential of base editing in multiplexing with tandem gRNA cassettes. Based on the metabolic pathways (Fig. 2A), we speculated that the disruption of two glycogen degradation pathways may further and stably improve the accumulation of glycogen. Therefore, we generated the working plasmid pSY-06 with gRNA-03 (targeting *glgP*) and gRNA-04 (targeting *glgX*) to inactivate *glgP* and *glgX* simultaneously (Fig. 3A and 3B). By applying pSY-06, we successfully introduced premature stop codons in *glgP* and *glgX* at the first round of selection with an editing efficiency of 80.77%. However, all selected colonies showed mixed sequencing signals at desired loci, forcing us to re-streak the edited cells for pure populations. After one more round of segregation, we obtained two edited strains, SY7 (*glgP* Gln84* and *glgX* Gln120*) and SY8 (*glgP* Gln84* and *glgX* Gln120* with a bystander missense mutation at His121 of *glgX*) (Fig. 3C).

**Fig. 3.**
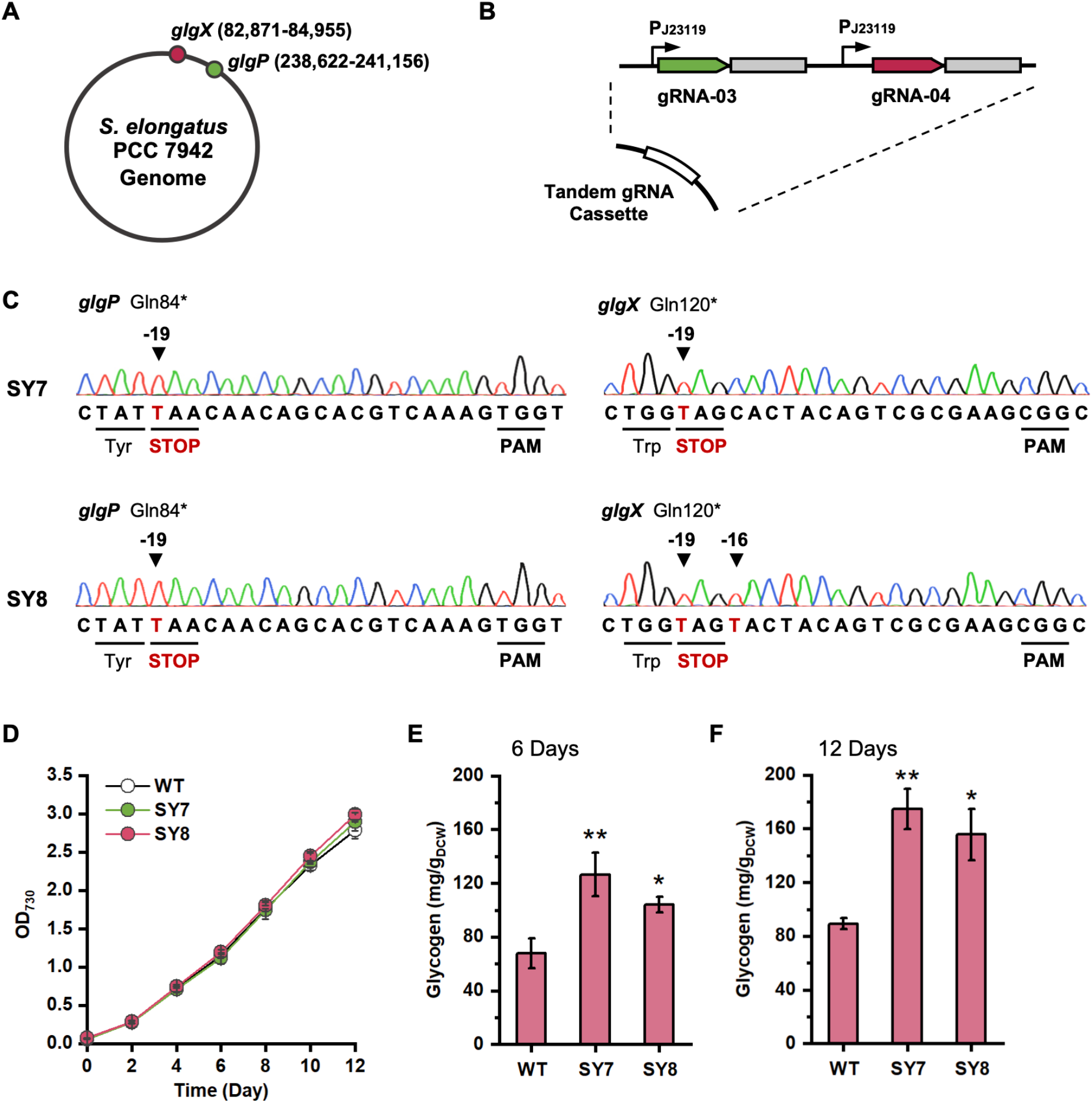
Multiplex base editing with tandem gRNAs. **(A)** The locations of *glgP* (238,622-241,156) and *glgX* (82,871-84,955) in the genome of *S. elongatus* PCC 7942. **(B)** The scheme of the tandem gRNA cassette with gRNA-03 and gRNA-04. **(C)** The sequencing results of the edited strains in the genes *glgP* and *glgX*. The edited loci were highlighted by the black arrow and the positions were counted from the PAM, regarding the first base on the left side of a PAM as position -1. **(D)** Growth profiles for the edited strains (SY7 and SY8) and the wild-type strain via the measurement of OD_730_ every 24 h. **(E)** Glycogen contents measured on day 6. **(F)** Glycogen contents measured on day 12. The error bars represent the standard deviations of three independent experiments. The differences in glycogen contents between the edited strains and the wild type were verified by *t*-test (* for P < 0.05 and ** for P < 0.01).

Then, we cultivated SY7, SY8 and the wild-type strain, and measured the glycogen contents. The SY7 and SY8 displayed similar growth profiles to that of the wild-type strain (Fig. 3D), indicating that the simultaneous inactivation of *glgP* and *glgC* would not influence the growth of cyanobacteria. We observed that the glycogen content in SY7 increased by 86.45% compared with that of the wild-type strain (67.86 ± 11.29 mg/gDCW) on day 6 (Fig. 3E), and was up to 174.69 ±14.89 mg/gDCW on day 12, achieving a 95.72% increase (Fig. 3F). SY8 also displayed higher glycogen contents, though slightly lower than SY7, on day 6 and day 12, which increased by 53.61% and 74.51% compared to that of the wild-type strain. Thus, the simultaneous inactivation of *glgP* and *glgX*, as expected, led to a significant increase in glycogen content from 26.70% to 95.72% on day 12 compared to the single inactivation of *glgP*. Overall, we demonstrated that our base-editing system is able to perform multiplex genome editing in cyanobacteria by inactivating two target genes at once, which is the first demonstration of multiplex genome editing in *S. elongatus*. Although an extra segregation step may be required, multiplex base editing saves a significant amount of time and labor to engineer cyanobacteria, which will be more beneficial in extensive metabolism reprogramming.

### Whole-genome sequencing reveals the off-target events

As dCas9 was employed for targeting and deamination might occur at untargeted loci, off-target events are possible when using the base-editing system. We carried out whole-genome sequencing to evaluate the off-target events of our base-editing systems. All the generated strains were whole-genome sequenced for the identification of single-nucleotide variations (SNVs). Considering the accumulation of SNVs with long-term cultivation, we also sequenced the wild-type strain in our lab for the background level of SNVs compared with the reference sequence of *S. elongatus* 7942 (NC_007604). The sequencing results showed that the wild-type *S. elongatus* PCC 7942 that we used has 16 SNVs, and these existing SNVs were excluded from the observed SNVs in the edited strains (Fig. 4A and Table S3).

**Fig. 4.**
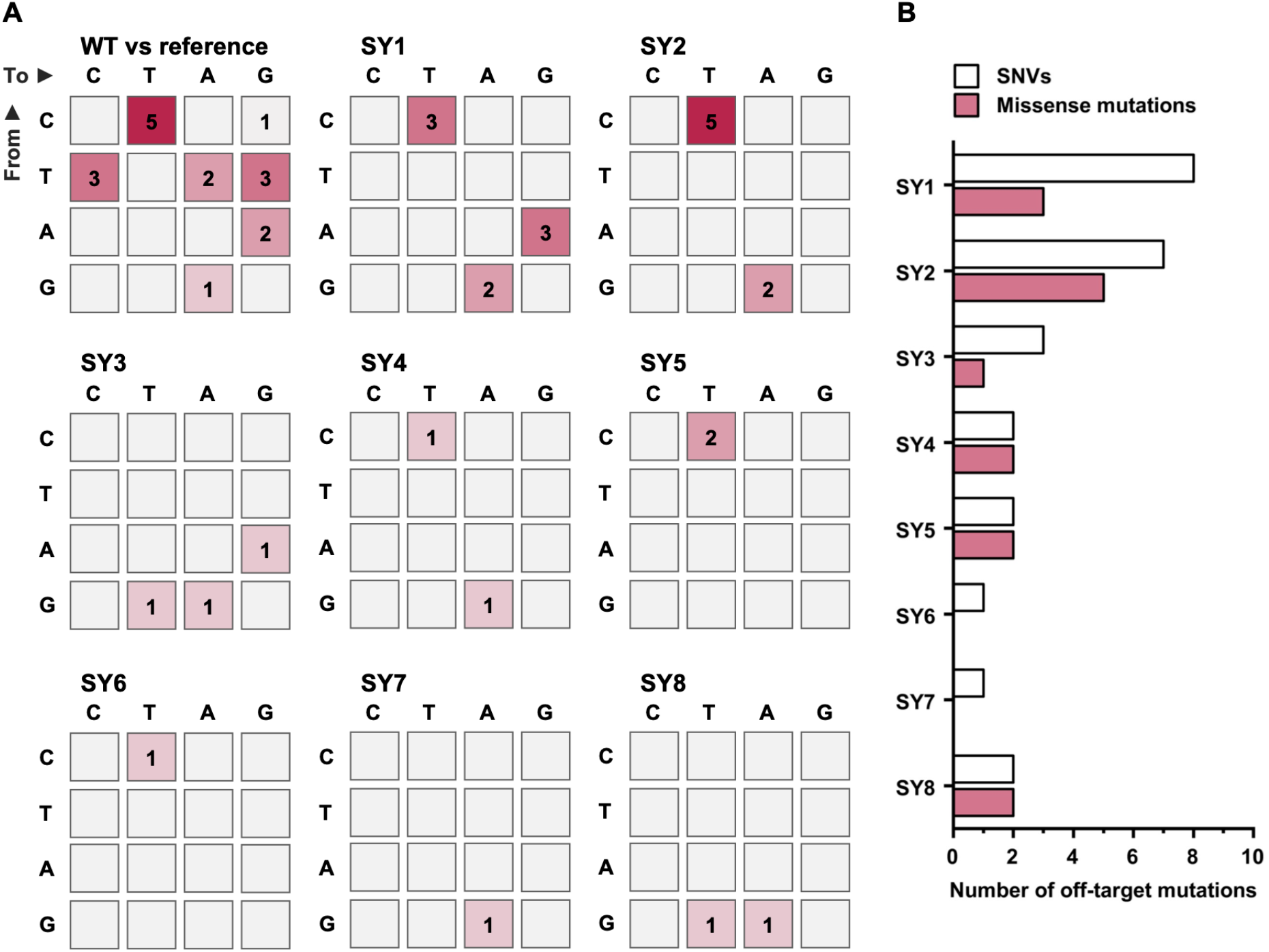
Off-target evaluation of the edited *S. elongatus* strains. **(A)** SNVs in the wild-type strain against the reference sequence of *S. elongatus* 7942 (NC_007604), and the off-target SNVs in the edited strains, where the sequenced SNVs in the wild type were excluded. The number in a particular cell indicates the type and quantity of SNVs. **(B)** Off-target SNVs and the resulting missense mutations in the edited strains.

First, we identified all designed on-target editing in the assigned loci in all edited strains. Then, we observed a small number of off-target SNVs in the edited strains, including 8 and 7 off-target SNVs in SY1 and SY2, which were slightly higher than other strains with less than 3 off-target SNVs. Meanwhile, only 1 and 2 off-target SNVs were found in SY7 and SY8 those were generated by the multiplex base editing (Fig. 4A). Among all these off-target events, the C-to-T and G-to-A substitutions took up the majority of mutations. They accounted for 33.33% in SY3, 50% in SY8, 62.5% in SY1, and 100% in other edited strains (Fig. 4A). Then, we investigated all off-targets loci to identify the correlation between off-target sequence and the gRNAs. Only one off-target SNV was found in SY4, showing a similar sequence to the corresponding gRNA-03 (Fig. S4), which indicated that most of the off-target events were caused by deamination rather than the CRISPR-Cas system. The results were in line with the results of applying the cytidine base editor in streptomyces, which indicated that most off-target SNVs resulted from deamination (28). Similar observations were also reported in, for instance, *Agrobacterium* strains and mouse embryos (29, 38). Considering that SNVs may generate missense or nonsense mutations which could possibly lead to undesired phenotypical variations, we checked all the off-target SNVs located in the coding sequences to characterize the mutations. Except for SY2 which has 5 off-target missense mutations, all strains showed less than three missense mutations among the off-target events, and, to be noticed, SY6 and SY7 exhibited no missense mutations (Fig. 4B). Moreover, no nonsense mutations at the off-target loci have been identified in all edited strains. Therefore, the off-target events generated by our base-editing system are seldom and the probability of unexpected phenotypes driven by the resulting mutations is low. However, off-target events do occur when using base editing, and comprehensive analysis of off-target events should be performed when necessary.

## Discussion

Cyanobacteria is an important microbial chassis for biological science discovery (39-41) and sustainable biotechnology (2, 42, 43). They can utilize CO_2_ from the atmosphere at the concentration of 400 parts per million, which is high enough to jeopardize the global climate but too low for biotechnology. To better understand and utilize cyanobacteria, methods for genetic perturbation in cyanobacteria have been invented almost four decades ago (6), and the first report of engineering cyanobacteria for ethanol production can be traced back to 1999 (44). Nowadays, cyanobacteria have been genetically reprogrammed to produce a variety of products (2, 4), and they have also been proposed as possible food suppliers not only on Earth but also on Mars (45, 46).

Despite the pioneering histories, the development of genetic tools for cyanobacteria, especially the CRISPR-Cas-based genome editing, seems to lag behind in the era of synthetic biology. We conclude two main obstacles slowing down the applications of CRISPR-Cas systems in cyanobacteria: 1) the toxicity of CRISPR-Cas systems and 2) the lack of self-replicating plasmids as carriers. To build a base-editing system, we also need to overcome these obstacles. First, we employed dCas9 as the CRISPR effector, which exhibits low toxicity and has been promoting inspiring progress in cyanobacteria (37, 47). The *lacI*-P_trc_ inducible system was utilized to drive the editing module for reducing the toxicity furthermore. Then, we chose pAM4787 as the backbone plasmid, which has been demonstrated self-replicable in *S. elongatus* PCC 9742 and *Anabaena* PCC 7120 (34). One extra advantage of pAM4787 is the lack of the stability module, the *pmaAB* or TA1 toxin/antitoxin system, which makes it relatively unstable in *S. elongatus* (34). This feature is highly favorable as a vector for genome editing tools, because it can carry the genetic tool well and can be cured with ease when the editing was complete. Given these rational designs, we successfully tailored and adapted base editing for cyanobacterium *S. elongatus*, and achieved genome editing at a single-nucleotide resolution, providing not only a new genome editing tool but also a paradigm for developing CRISPR-Cas-based genetic systems in cyanobacteria.

Besides the promising efficiency and precision of the base-editing systems, we found that clean editing can be identified at the first round of selection without extra segregation steps, though we cannot select clean editing in every editing experiment. The mixed populations of edited strains seem to be a common issue for base editing, which have been observed in different microbes, such as *C. ljungdahlii, Clostridium beijerinckii*, and *Agrobacterium* spp. (29, 31, 36). Moreover, we achieved, to our best knowledge, the first multiplex genome editing in the species of *Synechococcus*, and two distant loci with completely different protospacers were edited simultaneously (Fig. 3). Though a segregation step was required to obtain the pure population of the edited strains, the multiplex genome editing saved at least one whole round of editing experiments, including transformation, induction, selection, segregation (optional) and plasmid curing. Considering the relatively long doubling time of *S. elongatus* (6 – 8 h) compared to heterotrophs (48), our method exhibited a great potential to advance and accelerate the synthetic biology of cyanobacteria considerably with minimal time and labor.

Editing one nucleotide in a genome is powerful enough to reprogram the metabolism of a microbe. For instance, a previous work applied base editing in *C. ljunghdalii* and inactivated the aldehyde:ferredoxin oxidoreductase to improve acetate production from CO_2_ and H_2_ and to eliminate the ethanol accumulation (31). The produced acetate can be utilized by heterotrophic hosts for bioproduction (49). In the present study, we employed base editing to manipulate the glycogen metabolic pathway in cyanobacteria via introducing premature STOP codons into the relevant genes. The inactivation of *glgC* successfully reduced the accumulation of glycogen (Fig. 2D and 2E), and the carbon flux could be rediverted for producing other chemicals, such as isobutanol and succinate (10, 11). Meanwhile, the inactivation of *glgP* and the simultaneous inactivation of *glgP* and *glgX* significantly improved the glycogen content, especially the latter which generated a 95.72% increase compared to the wild-type strain. As glycogen is a polymer of glucose, it can be processed as food products, making the glycogen over-accumulating strain (e.g., SY7) a potential food supplier that uses atmospheric CO_2_ as the carbon source. It is reported by the Intergovernmental Panel on Climate Change (IPCC) that the CO_2_ emissions from the global food system attribute to 10.8 to 19.1 Gt CO_2_ -equivalent emissions per year, corresponding to 21% to 37% of the overall anthropogenic emissions of CO_2_ (50). Thus, a cyanobacterium-based food producer, as an alternative or complementary, may reduce the CO_2_ emission from conventional agriculture and support the ever-growing human population.

In summary, we present here a base-editing tool for cyanobacterium *S. elongatus* and achieved precise multiplex genome editing. Despite the successes, we want to point out the drawbacks of our base-editing system. First, the system still cannot achieve clean editing in every attempt immediately at the first round of selection, suggesting that an update or optimization of the editing module is required. Then, we cannot fully eliminate the off-target events, and comprehensive analysis of the off-target events for the edited strains may be necessary. Finally, single-nucleotide variations generated by base editing cannot fulfill all the tasks in reprogramming a microbe, such as gene insertion and deletion. Novel genetic tools for cyanobacteria are still in demand. In this endeavor, our base-editing tool will be a paradigm for overcoming the challenges in engineering cyanobacteria and for inventing new genome editing tools. We believe that our work will accelerate the synthetic biology of cyanobacteria and drive more carbon-negative innovations to alleviate global climate change.

## Methods

### Strains and media

All the strains used in this study are listed in Table S4. *Escherichia coli* DH5α was used for molecular cloning and was cultivated at 37 °C in LB medium (containing per liter: 10 g tryptone, 5 g yeast extract, and 10 g NaCl) or on solid LB agar plates (1.5%) supplemented with spectinomycin (60 μg/mL) or ampicillin (100 μg/mL) when required. Cyanobacterium *S. elongatus* PCC 7942 (ATCC 33912) was used as the wild-type strain. BG-11 and nitrate-depleted BG-11 media were employed for the cultivation of *S. elongatus* strains. Both spectinomycin and streptomycin were added to the medium at the concentration of 2 μg/mL when necessary. Cultures were grown in a climatic chamber at 30 °C under constant illumination (2000 – 3000 lux). The growth of *S. elongatus* strains was measured by the optical density at 730 nm (OD_730_). One liter of *S. elongatus* at OD_730_ 1.0 equals 0.334 g dry-cell.

### Plasmid construction

All the plasmids used in this study are summarized in Table S5. pSY was constructed in two steps. First, the editing module consisting of *dcas9, PmCDA1, ugi* and the LVA tag from pScI_dCas9-CDA-UL was cloned into pAM2991 (51), generating the editing module with the inducible system *lacI*-P_trc_. Then, the cassette was amplified and cloned into pAM4787, thus creating pSY. The gRNA cassettes were generated by changing the 20 bp of spacers via inverse PCR with pTemplate as the template. Finally, a modularized method was developed to generate the working plasmid serials. For instance, the pSY-01 was built by fusing the gRNA-01 cassette to pSY via DNA assembly methods. To generate pSY-06 for multiplex genome editing, the gRNA-04 was embedded into pSY-03, generating a tandem gRNA cassette. The PrimerSTAR Max DNA Polymerase (Takara Biotech.) was used for PCR, and the In-Fusion HD Cloning kit (Takara Biotech.) was used for DNA assembly. Plasmids were quantified by NanoDrop One (Thermo Fisher Scientific). All primers used in this study can be found in Table S6.

### Transformation and base editing

Transformation of *S. elongatus* PCC 7942 was performed following the protocol previously established (52). *S. elongatus* strains were growing in BG-11 medium until OD_730_ 0.7 - 0.9. Then, 15 mL of culture was harvested by centrifugation at 5,000 rpm for 10 min. The cells were washed and resuspended in 300 μL BG-11. After adding the working plasmids, the cells were cultivated in dark at 30 °C for 12 h before cultivating in fresh BG-11 medium for 3 days. The cells were then plated on selective BG-11 plates for the transformation of a plasmid not for genome editing. For base editing, two workflows were employed (Fig. 1A). For the direct-induction method, the cells after 3-d cultivation were plated on selective plates with IPTG as the inducer (1 mM). For the liquid-induction method, the cells after 3-d cultivation were induced with IPTG in the liquid medium for 16 – 24 h and then were plated on selective plates without an inducer. The editing results were determined via sequencing the target loci of randomly selected colonies. When necessary, one more round of segregation was performed to obtain the pure strain from the mixed populations by re-streaking the cells on selective BG-11 plates.

### Plasmid curing

To cure the working plasmids, the edited strain was cultivated in BG-11 medium without antibiotics for 7 – 10 days. Then the cells were streaked on BG-11 plate (no antibiotics) until colonies could be identified. The colonies were randomly chosen and analyzed by colony PCR to determine the loss of the working plasmids. After being tested by PCR, the colonies showing no signal for the working plasmid were cultivated again in BG-11 medium with and without antibiotics, respectively. The plasmid was regarded as cured only when 1) the colony PCR showed no signal for the plasmid and 2) the strain failed to grow in the media with antibiotics.

### Measurement of glycogen

The glycogen in *S. elongatus* PCC 7942 strains was extracted by a method modified from the previous report (53). *S. elongatus* was growing in the BG-11 medium at 30 °C under constant light at 100 rpm, and the cells were collected on day 6 and day 12 for analysis. The cells were washed and resuspended in 50 μL of sterilized water followed by adding 200 μL of 30% (w/v) KOH. The mixture was incubated at 95 °C for 2 h for lysis, and after cooling down, chilled ethanol was added and incubated at -20 °C overnight to precipitate glycogen. The glycogen was collected by centrifuge at 12,000 rpm, washed with 70% and 98% ethanol in succession, dried at 60 °C for 20 min, and resuspended in 100 μL of 100 mM sodium acetate (pH=5). Finally, the glycogen was digested to glucose by amyloglucosidase (5 U) at 60 °C for 2 h before analysis. The glucose was measured by High-Performance Liquid Chromatography (HPLC) (Shimadzu, Kyoto, Japan) using a ROA-Organic Acid H+ column (Phenomenex) and a reflective index detector (RID-20A, Shimadzu) at 50 °C. Sulfuric acid (0.005 mol/L) was used as the mobile phase at a flow rate of 0.6 mL/min.

### Whole-genome sequencing and analysis

Whole-genome sequencing was performed by Illumina sequencing to evaluate the off-target events in the edited *S. elongatus* strains. Briefly, around 30 mL of *S. elongatus* cells were harvested at the late-log phase (OD_730_ 0.8-1.0). First, the genome DNA was extracted and the genomic library was prepared by the TruSeq® DNA LT Sample Prep kit (Illumina Inc., USA). Then, the sequencing was carried out by an Illumina HiSeq Instrument. The results were cleaned, mapped and aligned using previously reported methods (54, 55). After being analyzed by the Qualimap software, the SNVs were determined by GATK4 (Haplotypecaller module) and Annovar (56, 57). The raw sequencing results will be found in the NCBI SRA upon acceptance.

## Acknowledgment

The authors thank the Core Facilities for Life and Environmental Sciences in Shandong University for the assistance with HPLC analysis. This work was supported by the National Natural Science Foundation of China (U20A20146), the Department of Science and Technology of Shandong Province (2022HWYQ-017), and the Qilu Young Scholar Program of Shandong University (to P.-F.X.).

## Supplementary data

Summary of strains, plasmids, primers and gRNA sequences; summary of the editing efficiency; summary of off-target events; and figures supporting the main results.

