## supplementary data for "Base Editing for Reprogramming Cyanobacterium *Synechococcus elongatus*"

**Table S1.** gRNA sequences used in this study.

| gRNA | Target | Strand [a] | PAM | Protospacer |
|---|---|---|---|---|
| gRNA-01 | *nblA* | C | AGG | TCTGCAGAAGTACCGTCAGC |
| gRNA-02 | *nblA* | C | CGG | AGCAGGTGCGTGACATCTCA |
| gRNA-03 | *glgP* | C | TGG | TCAACAACAGCACGTCAAAG |
| gRNA-04 | *glgX* | C | CGG | GCAGCACTACAGTCGCGAAG |
| gRNA-05 | *glgC* | C | CGG | GTTCCAAGGCACCGCCGATG |

[a] C stands for coding strand.

**Table S2.** Summary of sequencing results of base editing with pSY-01.[a]

| Induction methods | Direct induction [b] | Liquid induction [c] |
|---|---|---|
| Mixed edits | 20 | 23 |
| Clean edits | 8 | 10 |
| All edits | 28 | 33 |
| Overall editing efficiency [d] | 73.41 ± 13.27% | 86.51 ± 5.50% |
| Efficiency of clean editing [e] | 21.03 ± 12.50% | 26.98 ± 11.00% |
| Clean edits / All edits | 27.19 ± 12.86% | 31.79 ± 14.21% |

[a] Data were summarized based on 38 colonies screened from 3 independent rounds of editing experiments.

[b] Direct induction: transformants are induced while plating.

[c] Liquid induction: transformants are induced in liquid medium before plating.

[d] Overall editing efficiency was the percentage of the number of all edits, including clean edits and mixed edits, divided by the number of picked colonies in total.

[e] Efficiency of clean editing was the percentage of the number of clean edits divided by the number of picked colonies.

**Table S3.** The SNVs revealed by whole-genome sequencing.[a]

| Strains | Alteration | Gene | Position | Mutation type |
|---|---|---|---|---|
| Wild type | C to T | RS05015 | 985620 | missense mutation |
| | C to T | RS07925 | 1611805 | missense mutation |
| | G to A | RS10715 | 2195578 | nonsense mutation |
| SY1 | A to G | RS00245 | 49811 | silent mutation |
| | C to T | RS01325 | 256425 | missense mutation |
| | T to C | RS05015 | 985620 | missense mutation |
| | A to G | - | 1538363 | - |
| | T to C | RS07925 | 1611805 | missense mutation |
| | G to A | RS10275 | 2093794 | silent mutation |
| | A to G | RS10715 | 2194832 | silent mutation |
| | A to G | RS10715 | 2195578 | silent mutation |
| SY2 | G to A | RS00115 | 22723 | silent mutation |
| | C to T | RS03870 | 742224 | missense mutation |
| | C to T | RS09200 | 1883810 | missense mutation |
| | C to T | RS09220 | 1890576 | missense mutation |
| | C to T | RS10210 | 2082899 | missense mutation |
| | C to T | RS11865 | 2401832 | missense mutation |
| | G to A | RS12040 | 2438130 | silent mutation |
| SY3 | A to G | RS00245 | 49811 | silent mutation |
| | G to A | RS00695 | 139119 | silent mutation |
| | G to T | RS01245 | 238859 | missense mutation |
| SY4 | C to T | RS00010 | 2061 | missense mutation |
| | G to A | RS01470 | 284423 | missense mutation |
| SY5 | C to T | RS01440 | 280251 | missense mutation |
| | C to T | RS02685 | 513193 | missense mutation |
| SY6 | C to T | RS09815 | 2008471 | silent mutation |
| SY7 | G to A | RS00290 | 60438 | silent mutation |
| SY8 | G to A | RS01115 | 220611 | missense mutation |
| | G to T | RS01245 | 238859 | missense mutation |

[a] SNVs in the wild-type strain in our lab were identified by comparing with the reference genome (NC_007604), and they were excluded when determining the off-target SNVs in the edited strains.

**Table S4.** Strains used in this study.

| Name | Features | Source |
|---|---|---|
| *S. elongatus* PCC 7942 | Wild type | ATCC33912 |
| SY1 | Wild type, *nblA* Gln19* | This study |
| SY2 | Wild type, *nblA* Gln19*, silent mutation at Leu18 | This study |
| SY3 | Wild type, *nblA* Gln24* | This study |
| SY4 | Wild type, *glgP* Gln84* | This study |
| SY5 | Wild type, *glgX* Gln120* | This study |
| SY6 | Wild type, *glgC* Gln99* | This study |
| SY7 | Wild type, *glgP* Gln84*, *glgX* Gln120* | This study |
| SY8 | Wild type, *glgP* Gln84*, *glgX* Gln120* and His121Tyr | This study |

**Table S5.** Plasmids used in this study.

| Name | Features | Source |
|---|---|---|
| pAM2991 | *colE1* ori, *lacI*-$P_{trc}$ | Addgene #40248, (1) |
| pAM4787 | *colE1* ori, partial sequence from pANS | Addgene #120088, (2) |
| pScI_dCas9-CDA-UL | *pSC101* ori*,* *dcas9*, *PmCDA1*, *ugi* | Addgene #108551, (3) |
| pTemplate | *pUC* ori, $P_{J23119}$-gRNA (no spacer) | Lab stock |
| pSY | pAM4787, *lacI*, $P_{trc}$-*dcas9*-*PmCDA1*-*ugi* | This study |
| pTemplate-01 | pTemplate, gRNA-01 | This study |
| pTemplate-02 | pTemplate, gRNA-02 | This study |
| pTemplate-03 | pTemplate, gRNA-03 | This study |
| pTemplate-04 | pTemplate, gRNA-04 | This study |
| pTemplate-05 | pTemplate, gRNA-05 | This study |
| pSY-01 | pSY, gRNA-01 | This study |
| pSY-02 | pSY, gRNA-02 | This study |
| pSY-03 | pSY, gRNA-03 | This study |
| pSY-04 | pSY, gRNA-04 | This study |
| pSY-05 | pSY, gRNA-05 | This study |
| pSY-06 | pSY, gRNA-03, gRNA-04 | This study |

**Table S6.** Primers used in this study.

| Primer | Sequence |
|---|---|
| Primers for In-Fusion DNA assembly | |
| XIA-WSY-01 | cacacaggaaacagaccatgatggataagaaatactcaatagg |
| XIA-WSY-02 | agttccctactctcgcatggttatgcaaccagtcctagca |
| XIA-WSY-03 | cctattgagtatttcttatccatcatggtctgtttcctgtgtg |
| XIA-WSY-04 | tgctaggactggttgcataaccatgcgagagtagggaact |
| XIA-WSY-05 | cggataccggtttcgaattggcgaggcagcagatcaattc |
| XIA-WSY-06 | tgcccagctcggtctagattgcttcgcaacgttcaa |
| XIA-WSY-07 | gaattgatctgctgcctcgccaattcgaaaccggtatccg |
| XIA-WSY-08 | ttgaacgttgcgaagcaatctagaccgagctgggca |
| XIA-WSY-11 | GACagagacgcaagacacttggccaatgctcgagcgctt |
| XIA-WSY-12 | ggagtcactgccaaccgagacgcatgtgcttgggtgctgcgat |
| XIA-WSY-13 | aagcgctcgagcattggccaagtgtcttgcgtctctGTC |
| XIA-WSY-14 | atcgcagcacccaagcacatgcgtctcggttggcagtgactcc |
| XIA-WSY-15 | gaaggcttgcgatggttt |
| XIA-WSY-56 | gatggccagcgttccactatagtcgaaccacgcaatgc |
| XIA-WSY-57 | ttgcgatggtttgcggctaaaggaagcggcacacaggaaa |
| XIA-WSY-58 | tttcctgtgtgccgcttcctttagccgcaaaccatcgcaa |
| XIA-WSY-59 | gcattgcgtggttcgactatagtggaacgctggccatc |
| Primers for Inverse-PCR | |
| XIA-WSY-16 | tctgcagaagtaccgtcagcgttttagagctagaaatagc |
| XIA-WSY-17 | gctgacggtacttctgcagagctagcattatacctaggac |
| XIA-WSY-36 | tcaacaacagcacgtcaaaggttttagagctagaaatagc |
| XIA-WSY-37 | ctttgacgtgctgttgttgagctagcattatacctaggac |
| XIA-WSY-44 | agcaggtgcgtgacatctcagttttagagctagaaatagc |
| XIA-WSY-45 | tgagatgtcacgcacctgctgctagcattatacctaggac |
| XIA-WSY-52 | gttccaaggcaccgccgatggttttagagctagaaatagc |
| XIA-WSY-53 | catcggcggtgccttggaacgctagcattatacctaggac |
| XIA-WSY-54 | gcagcactacagtcgcgaaggttttagagctagaaatagc |
| XIA-WSY-55 | cttcgcgactgtagtgctgcgctagcattatacctaggac |
| Primers for colony PCR and sequencing | |
| XIA-WSY-21 | aatgctgctgcctctacttc |
| XIA-WSY-22 | gtgtaccagttgccaagcca |

| | |
|---|---|
| XIA-WSY-23 | cgagatagcagtattgacgg |
| XIA-WSY-42 | cgagttccatagcgttaagg |
| XIA-WSY-43 | gatcgccgaagtatcgactc |
| XIA-WSY-46 | tcatagcctttggactgaag |
| XIA-WSY-47 | agtagtcgttgaacagaaag |
| XIA-WSY-48 | gtagggtaagaaatcgcaaa |
| XIA-WSY-49 | gcgccagttgatttcgtaga |
| XIA-WSY-50 | ggatctgattcgttatatgc |
| XIA-WSY-51 | ctactacagccagggcaaaa |
| XIA-WSY-60 | tctgtgaggatgaacggact |
| XIA-WSY-61 | ctgccatgactgtttcatcc |
| XIA-WSY-62 | cctgttcatccagcagatag |
| XIA-WSY-63 | cacctctttctactggcatg |

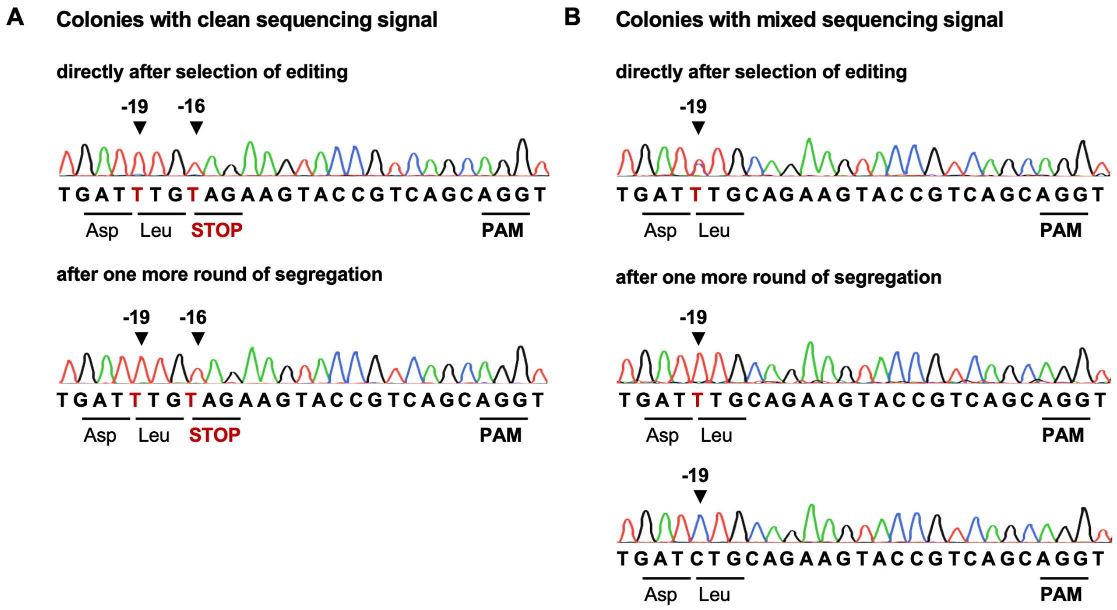


**Fig. S1. Sequencing results of edited colonies in *nblA* after one more round of segregation.** (A) The sequencing results of clean edited colonies immediately after editing and after an extra round of segregation. (B) The sequencing results of colonies with mixed signals in *nblA* after editing and after one more round of segregation. The sequenced colonies were edited with pSY-01 (gRNA-01).

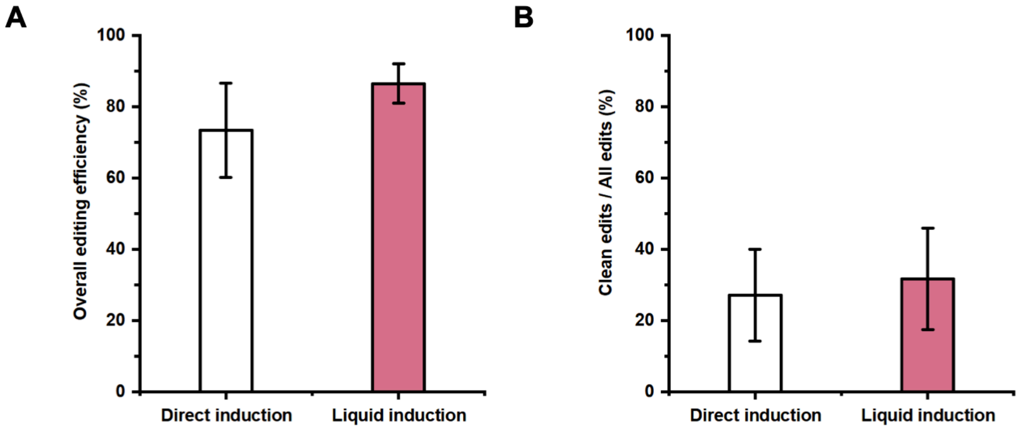


**Fig. S2. The comparison of editing efficiency between two induction methods.** (A) The comparison of overall editing efficiency, the percentage of the number of all edits, including clean edits and mixed edits, divided by the number of picked colonies. (B) The comparison of the ratios between the number of clean edits and all edits. The editing was performed with pSY-01 and the data were collected from three independent editing experiments, and error bars represent the standard deviations.

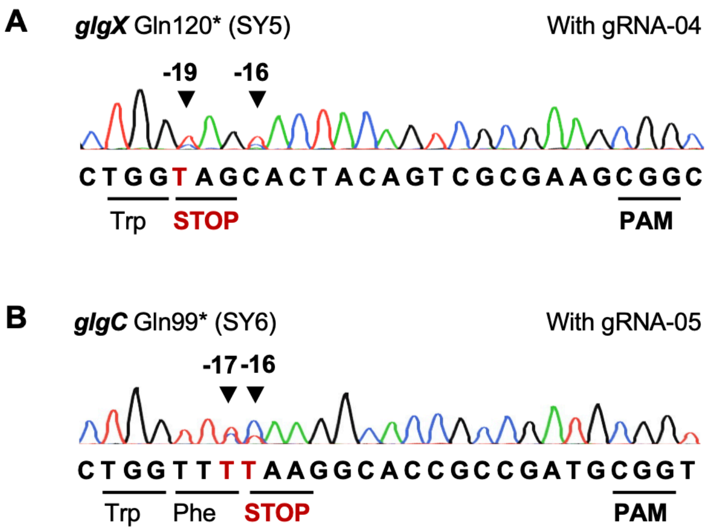


**Fig. S3. Mixed sequencing signals and bystander editing.** (A) Mixed sequencing signals on two loci of *glgX* after being edited with pSY-04 (gRNA-04), and the bystander editing loci located at position -16, while position -19 was the target locus. (B) Mixed sequencing signals on two loci of *glgC* after being edited with pSY-05 (gRNA-05), and the bystander editing loci located at position -17. The edited loci were highlighted by the black arrow and the positions were counted from the PAM, and the first base on the left side of a PAM was regarded as position -1.

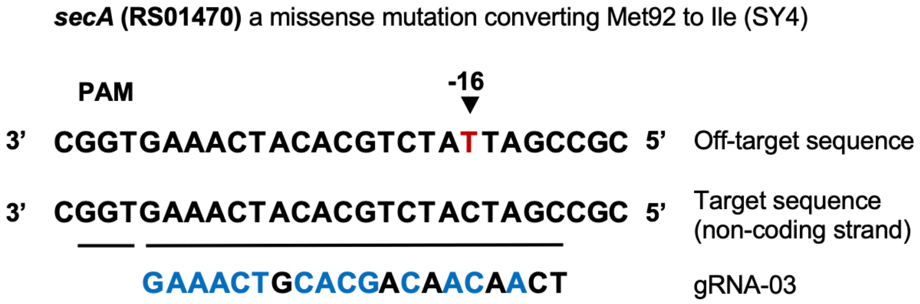


**Fig. S4. The sequences of the target sequence, off-target sequence and gRNA-03.** The same bases in gRNA-03 were marked blue compared with the probable spacer of the off-target SNV in SY4. The edited loci were highlighted by the black arrow and marked red. The positions were counted from the PAM, and the first base on the left side of a PAM was regarded as position -1.
